# Complex impact of stimulus envelope on motor synchronization to sound

**DOI:** 10.1101/2024.08.06.606724

**Authors:** Yue Sun, Georgios Michalareas, Oded Ghitza, David Poeppel

## Abstract

The human brain tracks temporal regularities in acoustic signals faithfully. Recent neuroimaging studies have shown complex modulations of synchronized neural activities to the shape of stimulus envelopes. How to connect neural responses to different envelope shapes with listeners’ perceptual ability to synchronize to acoustic rhythms requires further characterization. Here we examine participants’ motor and sensory synchronization to noise stimuli with periodic amplitude modulations (AM). We used three envelope shapes that varied in the sharpness of amplitude onset. In a synchronous motor finger-tapping task, we show that participants more consistently align their taps to the same phase of stimulus envelope when listening to stimuli with sharp onsets than to those with gradual onsets. This effect is replicated in a sensory synchronization task, suggesting a sensory basis for the facilitated phase alignment to sharp-onset stimuli. Surprisingly, despite less consistent tap alignments to the envelope of gradual-onset stimuli, participants are equally effective in extracting the rate of amplitude modulation from both sharp and gradual-onset stimuli, and they tapped consistently at that rate alongside the acoustic input. This result demonstrates that robust tracking of the rate of acoustic periodicity is achievable without the presence of sharp acoustic edges or consistent phase alignment to stimulus envelope. Our findings are consistent with assuming distinct processes for phase and rate tracking during sensorimotor synchronization. These processes may be underpinned by different neural mechanisms whose relative strengths are modulated by specific temporal dynamics of stimulus envelope characteristics.

## Introduction

Temporal regularity is a fundamental feature of our acoustic environment, including in speech and music (Ding et al., 2017; Varnet et al., 2017). The human auditory system detects temporal regularities by tracking fluctuations in the amplitude envelope of acoustic signals (e.g., Doelling et al., 2014). Neural synchronization to stimulus envelope has been demonstrated using a wide range of stimuli, from click-type stimuli with sharp amplitude onsets (Lakatos et al., 2013; ten Oever et al., 2017) to noise/musical sounds with gradual amplitude onsets (Doelling et al., 2019; Henry et al., 2014).

Recent neurophysiological studies suggest that the shape of stimulus envelope may impact the strength of stimulus-brain synchronization (e.g., Doelling et al., 2019; Oganian & Chang, 2019). For instance, Irsik et al. (2021) recorded participants’ electroencephalographic (EEG) responses when they listened to periodic sequences of noise stimuli with either sharp or gradual onsets. They showed that both envelope shapes can induce periodic and synchronized neural activities from auditory cortex, albeit with varied responses to stimuli from each envelope type. On the one hand, the authors observed that sharp-onset stimuli elicit stronger auditory event-related potentials (ERPs) than gradual-onset stimuli, in line with previous studies (Doelling et al., 2019; Thomson et al., 2009). Evoked responses to salient acoustic landmarks have been proposed to underpin efficient tracking of temporal regularity in acoustic input (Oganian & Chang, 2019; Zou et al., 2021). On the other hand, gradual-onset stimuli evoke smoother, more sinusoidal amplitude patterns in neural activity, which also exhibit higher inter-trial coherence (Irsik et al. 2021). The authors interpreted this finding as stronger neural synchronization to the repetition rate of stimuli. This interpretation resonates with a previous study showing more consistent phase alignment between slow cortical activity and stimulus envelope when participants listened to sequences with gradual-onset stimuli than those with sharp-onsets (Doelling et al., 2019).

The neuroimaging findings point to an interesting dichotomy: despite the significant differences in the neural evoked responses to sharp versus gradual envelope onsets, the brain can successfully track the rhythmic structure in the acoustic input with both types of envelopes. This pattern motivates an investigation into the higher-order characteristics that underlie such robust brain-stimulus synchronization. Here, in two behavioral paradigms and a suggested model, we examine how the stimulus envelope influences listeners’ ability to behaviorally synchronize to acoustic input. We aim to build connections between variable neural responses to acoustic envelopes, observed in previous neuroimaging studies, and higher-order aspects of human listeners sensory and motor synchronization to auditory rhythmicity.

Participants performed in a sensorimotor synchronization task (Repp, 2005; Repp & Su, 2013), which requires synchronous finger-tapping to noise stimulus sequences with periodic amplitude modulations. We used three modulation envelopes (Figure 1A): damped (sharp onset, gradual offset), ramped (gradual onset, sharp offset) and symmetrical (gradual onset, gradual offset). All three envelopes have been commonly used in neurobiological studies of auditory processing in humans (Henry et al., 2014; Irsik et al., 2021) as well as in non-human animals (Herrmann et al., 2017; Liu & Wang, 2022). However, the behavioral synchronization to these envelopes is not well characterized.

**Figure 1.**
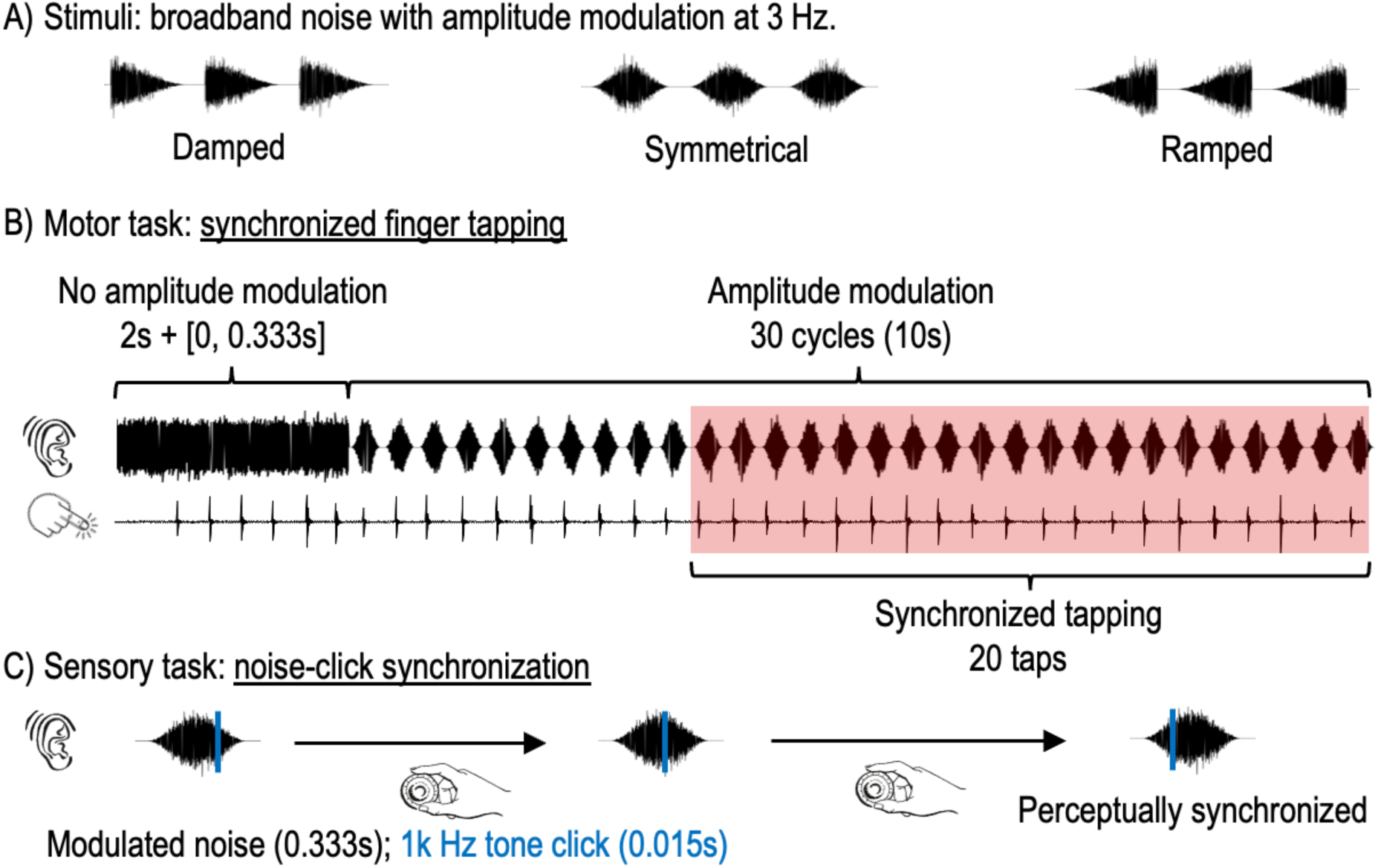
Stimuli and paradigm. A) Sample excerpts of acoustic stimuli with different envelope shapes. Each sample shows 1 second of acoustic signal which comprises three cycles of the amplitude modulated noise with the damped, symmetrical, and ramped envelopes (the modulation frequency is 3 Hz). B) Paradigm for the motor synchronization task. The schematic illustrates a single trial of the task. The top row depicts the acoustic stimulus, the bottom row indicates the timing of each finger tap from the participant in response to the acoustic input. The red area highlights the analysis time window during which participants are assumed to have reached synchronized tapping. The time window comprises the final 20 taps of each trial. C) Paradigm for the sensory synchronization task. During this task, participants are presented with the same noise sequences as in the motor synchronization task. Along with each cycle of the modulated noise, a 1-kHz tone click (blue line) is presented, with its initial location being randomly selected within the modulation time window. Both the modulated noise and tone click are presented diotically. Participants are instructed to change the location of the tone click using a dial until the click is perceived to be synchronized to the noise stimuli. Participants then press a button to confirm the final location of the click, marking the end of the trial.

We examine the impact of these envelopes on two aspects of participants’ synchronization performance: (i) the ability to continuously align their taps to the same part (or phase) of the repeated stimulus envelope and (ii) the ability to maintain their tapping rate at the modulation rate of the acoustic stimuli. These two aspects have been demonstrated to involve different cognitive processes, namely phase and rate tracking, that occur in parallel during motor synchronization to periodic acoustic signals (Jacoby & Repp, 2012). In addition to the motor synchronization task, we also conducted a sensory synchronization experiment to examine the sensory basis of the envelope effects revealed in the motor task. Finally, we describe a model to qualitatively account for the envelope effect observed in both sensory and motor experiments.

## Methods and Materials

### Participants

Twenty-four participants (14 females, 10 males; average age: 22.04, range 18 to 27) provided written informed consent to take part in the study and received monetary compensation for their participation. All participants were right-handed and reported normal hearing. The experimental procedure was approved by the Ethics Council of the Max-Planck Society (no. 2017_12).

### Experimental design

#### Stimuli

We used broadband (0-22050 Hz) Gaussian noise stimuli which were amplitude modulated (AM) at 3 Hz. Each modulation window (333 ms) was composed of an AM noise stimulus (273 ms) that was preceded and followed by a silence period of 30 ms (in total 60 ms of silence in each modulation window). We used three modulation envelopes: *damped*, *symmetrical* and *ramped*. The *damped* envelope corresponded to the descending half-cycle (peak to trough) of a sinusoidal function with a period of 273 × 2 = 546 ms (Figure 1A, left), such that noise stimuli with this envelope exhibited sharp (abrupt) amplitude onset and gradual amplitude offset. The *symmetrical* envelope corresponded to the full cycle (trough to trough) of a sinusoidal function with a period of 273 ms (Figure 1A, middle), such that noise stimuli with this envelope exhibited symmetrical amplitude rise and decay with the peak amplitude in the middle of the modulation window. The *ramped* envelope corresponded to the ascending half-cycle (trough to peak) of a sinusoidal function with a period of 273 × 2 = 546 ms (Figure 1A, right), such that that noise stimuli with this envelope exhibited gradual amplitude onset and sharp (abrupt) amplitude offset. The RMS of each modulation window was normalized across different stimulus envelope shapes.

#### Tasks

Each participant performed two behavioral tasks. The first is a sensorimotor synchronization task in which participants were asked to produce continuous finger taps in sync with the periodically presented AM noise stimuli (Figure 1B). The second task consisted of a sensory synchronization task in which the periodic AM noise sequence was simultaneously presented with a sequence of pure tone clicks that had the same presentation rate (Figure 1C). Participants were instructed to adjust the delay between the tone sequence and noise sequence until the two acoustic streams were perceived as in-sync with each other.

The sensory synchronization task was introduced to complement findings from the motor synchronization task. Due to the absence of motor output, the sensory synchronization task examined participants’ perception of the acoustic landmarks within the envelope of the noise stimuli that elicit the percept of synchronization with another acoustic stream. The same acoustic landmarks are assumed to serve as sensory references to guide the motor synchronization in the finger tapping task. Results from the sensory task would by hypothesis reveal how the envelope shape impacts the level of variability in the sensory processing of these landmarks, and thus highlight the sensory components of synchronization variability in participants’ performances in the motor task.

### Procedure

Participants were seated in a sound-proof booth in front of an LCD monitor to receive instructions and feedback. Auditory stimuli were generated using MATLAB (The MathWorks, Natick, MA, USA) at 44.1 kHz/16 bits, output by a high-quality interface (RME Fireface UCX) and presented to participants binaurally via electrodynamic headphones (Beyerdynamic DT770 PRO). The experiment was run using MATLAB Psychophysics Toolbox extensions (Brainard, 1997) on a Fujitsu Celsius M730 computer running Windows 7 (64 bit).

#### Motor synchronization task

Participants were instructed to lay their right forearm and hand on the desk and tap their right index finger on the desk surface during each trial next to a Schaller Oyster S/P contact microphone. The sound of their taps was collected by the microphone and recorded via the RME Fireface UCX soundcard.

In each trial, one periodic sequence of amplitude modulated (AM) broadband noise with a modulation rate at 3 Hz was presented to the participant. Each sequence contained 30 modulation cycles, leading to a duration of 10 seconds. The average output intensity of the modulated sequence was calibrated at 70 dB (A-weighted). In each trial, the 10-second AM noise sequence was always preceded by a short noise stimulus whose amplitude was fixed at the maximum level of the amplitude modulation (Figure 1B). The duration of this unmodulated noise stimulus varied randomly between 2 and 2.333 seconds across different trials.

Participants were instructed to start tapping their finger continuously after the onset of the unmodulated noise with any speed and rhythms they liked. Given that the duration of the unmodulated noise was variable across trials, having participants engage in free-style tapping alongside this noise stimulus assured a random alignment between participants’ taps and the stimulus envelope at the moment when the periodic amplitude modulation started. After the beginning of the amplitude modulation, participants’ task was to transition from free-style tapping to synchronized tapping to the periodic presented noise stimuli as fast as possible. Once synchronized tapping had been reached, they needed to maintain the synchronized tapping until the end of the sequence. After each trial, an inter-trial interval randomly distributed between 1 and 2 seconds occurred before the beginning of the next trial.

For this task, each participant performed a total of 90 trials, divided into 10 blocks. Each block contained 9 trials, 3 for each envelope shape. The trial order within each block was pseudo-randomized such that there was no repetition of the same envelope condition between any two adjacent trials. Each block lasted about 2 minutes, and participants were given a short break after each block. The experiment started with a practice session composed of 6 trials (two trials for each envelope condition), such that participants could familiarize themselves with the trial structure and their task, and find a comfortable arm and hand position for the experiment. Data from this session were excluded from the analysis.

#### Sensory synchronization task

On each trial, participants received a periodic sequence of AM noise stimuli with one of the three stimulus envelopes. Individual AM stimuli within each sequence were created with the same physical characteristics as in the motor synchronization task. Therefore, the AM noise sequence exhibited a 3 Hz modulation rate. Concurrently with the noise stimuli, a 1-kHz pure tone click was presented within each modulation window. The tone click had a 10-ms duration with 1-ms rise and decay time and it was presented with a signal-to-noise (SNR) level of 2 dB with respect to the AM noise stimuli. This SNR level was selected such that the tone stimulus can be perceived at all locations of the modulation window without being too intense. The noise stimuli and tone clicks were diotically presented to participants.

For each trial, the first tone click was presented at a random temporal position within the modulation window of the noise stimulus, which gave a random delay between the onset of the tone click and the onset the AM noise. Participants were instructed to adjust the temporal position of the tone click by rotating a physical dial (Griffin Technology PowerMate UBS knob) until they perceived the tone click as being synchronized with the AM noise stimuli. Turning the dial clockwise would move the tone click forward in time, while turning the dial anti-clockwise would move the tone backward. It is noteworthy that the tone click was moved within a circular space that covered the length of the modulation window of the noise stimuli. That is, moving the tone click beyond the right-end boundary of the modulation window would made it appear at the left-end boundary of that window, and vice versa. Since both acoustic streams (AM noise and tone click) were presented continuously, participants received the updated position of the tone click in real-time and could continuously assess the synchrony between the two streams. Upon their perceptual judgment of synchrony between the tone click and AM noise, participants were instructed to press a button on the keyboard to confirm the final position of the tone click. This confirmation would mark the end of the trial and stop the presentation of both stimuli. An inter-trial interval randomly distributed between 1 and 2 s occurred before the beginning of the next trial.

Participant also received 90 trials in total, divided into 10 blocks. Each block contained 9 trials, 3 for each envelope shape. The trial order within each block was pseudo-randomized such that there was no repetition of the same envelope condition between any two adjacent trials. For this task, since the duration of each trial was determined by participants, i.e. depending on how long they needed to reach the percept of synchrony, the duration of each block was variable. In average, the 10 blocks of the test lasted about 45 minutes. This experiment started with a practice session composed of 6 trials (two trials for each envelope condition). Data from this session were excluded from the analysis.

### Data analysis

#### Motor synchronization task

##### Detection of individual taps of each trial

The raw data of each trial consisted of a sound file that contained two channels. The first channel contained the sound of participants’ finger taps during the entire trial recorded by the external microphone. The second channel contained the sequence of noise stimuli that was internally recorded by the soundcard during the trial. In order to measure the onset timing of individual taps of each trial, we developed a procedure that enabled flexible amplitude thresholding to extract tap onsets (abrupt amplitude rises to a relatively high level) from the background noise floor and, occasionally, from other low amplitude artifactual sound that overlapped with the tap (e.g., sound caused by movements of the participants’ arm or hand against the desk surface). The trial recording as well as detected tap onsets were plotted for visual inspection, which allowed for further removal of sound artifacts that were falsely detected as taps.

##### Between-trial synchronization performance

Since we investigated participants’ performance in synchronized tapping to AM noise stimuli, we focused on participants’ final 20 taps of each trial (Figure 1B), during which participants were assumed to have reached synchronized tapping. From these taps, we derived several measurements for each trial which were later used to examine participants’ between-trial and within-trial synchronization performance. To assess participants’ between-trial performance, we examined the average location and variability across participants’ tap-stimulus alignments (TSAs) from different trials of each envelope condition. The essential measurement for these analyses was the TSA location for each trial (TSA_trial)_, which was computed by averaging the TSA across the 20 taps within the trial (Figure 2A). For this measurement, we first calculated the temporal delay (in ms) between the onset of each tap and the onset of the nearest modulation window. The cyclic presentation of AM noise stimuli led to a circular distribution of the TSAs. In order to more accurately estimate the average TSA location across the 20 taps of each trial, we converted the time delays (ms) into angles (radian) within a circular space that covered the length of the modulation window, and computed the circular mean of these angles, which gave the average TSA location of each trial (TSA_trial)_. We then averaged TSA_trial a_cross different trials of each envelope condition, which was used to assess participants’ overall TSA locations for each envelope shape (TSA_condition)_. To examine how consistently participants’ taps aligned to the same part of stimulus envelope across different trials of the same envelope condition, we computed the *circular standard deviation* (CSD) (Mardia, 1975) across TSA_trial f_rom different trials of each envelope condition (CSD_between-trial T_SA). Consistent tap-stimulus alignments between trials should lead to low CSD_between-trial T_SA.

**Figure 2.**
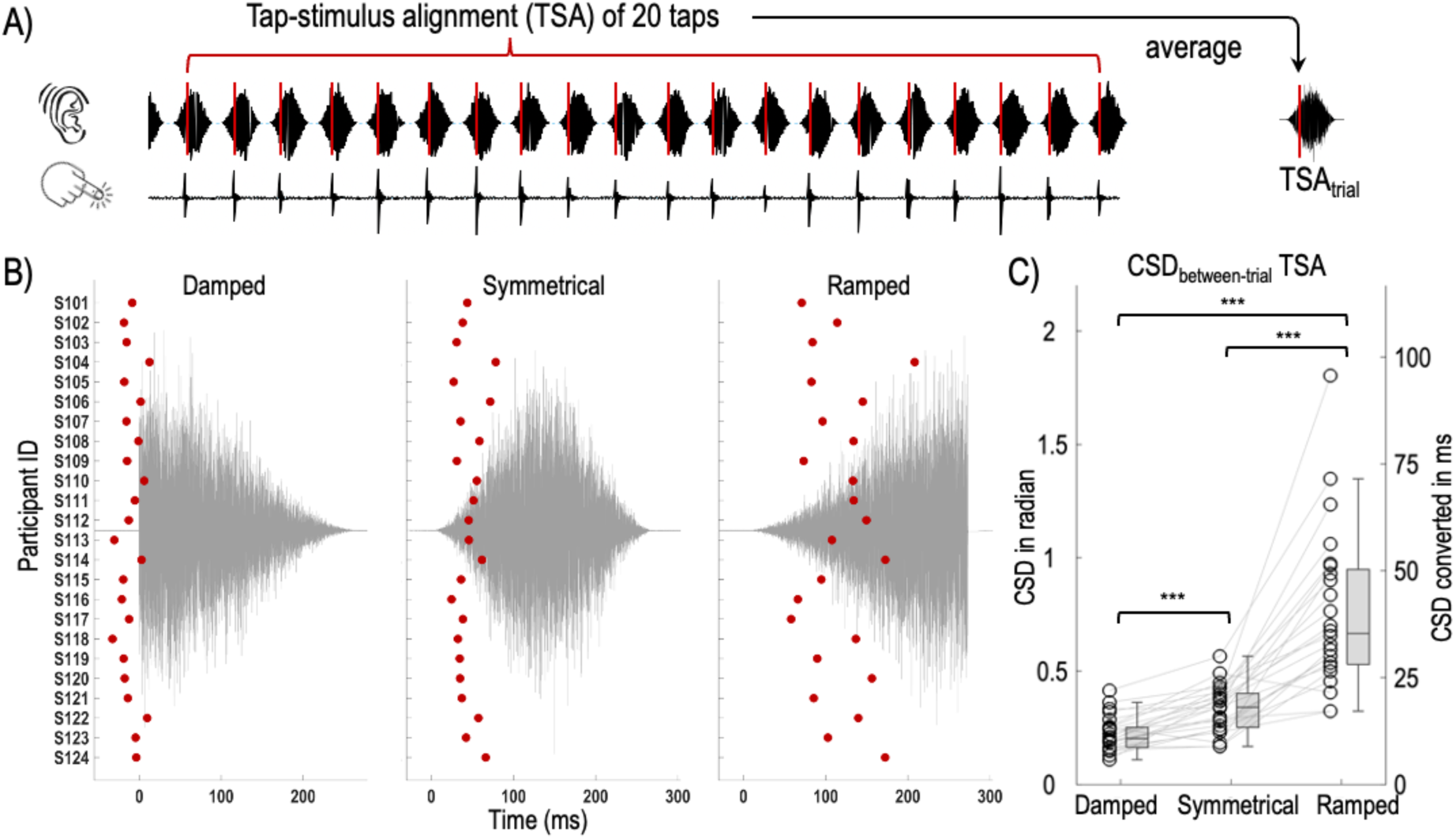
Participants’ between-trial performance in the motor synchronization task. A) Computation of the tap-stimulus alignment of each trial (TSA_trial_) by averaging the TSA across the 20 final taps of each trial. B) Average TSA of each stimulus envelope condition for individual participants (left: Damped stimuli; middle: Symmetrical stimuli; right: Ramped stimuli). In each graph, the grey curve indicates the wave form of an example stimulus with the corresponding stimulus envelope. X-axis represents time, with zero indicating the onset of the amplitude rise. The red circles indicate the condition-average of TSA (TSA_condition_) of each of the 24 participants. Y-axis shows participant ID, common across the three graphs. C) Between-trial variation of TSA of the three stimulus envelope conditions (CSD_between-trial_ TSA). For each envelope condition, circles indicate the average CSD_between-trial_ TSA of individual participants. The levels of CSD are shown in both radians (y-axis on the left side) and ms (converted from circular data) (y-axis on the right side). Asterisks mark statistical significance of pair-wise comparisons among envelope conditions (***: *p* < .001).

##### Within-trial synchronization performance

To assess participants’ within-trial performance, we examined their maintenance of the tapping rates and tap-stimulus alignments across the final 20 taps within each trial. To this end, we calculated the inter-tap-intervals (ITI) between every two adjacent taps across the 20 taps of each trial, which yielded 19 ITIs per trial (Figure 4A). The mean ITI of each trial thus indicated the tapping rate of the trial, which should be close to 333 ms upon synchronized tapping at 3 Hz. Meanwhile, the maintenance of the tapping rate within the trial would be reflected in the standard deviation across the 19 ITIs, which we referred to as SD_within-trial I_TI. Steady tapping rate within trials should lead to low SD_within-trial I_TI. To examine the maintenance of tap-stimulus alignments (TSA) within trials, we measured the TSAs of individual taps within each trial, first as time delay (ms) and then converted into angles (radian) within a circular space that covered the modulation window of noise stimuli (Figure 4A). To examine how consistently participants’ individual taps aligned to the same part of stimulus envelope within single trials, we computed the circular standard deviation (Mardia, 1972) across the 20 TSAs for each trial (CSD_within-trial T_SA). Consistent tap-stimulus alignment within trials should lead to low CSD_within-trial T_SA for each individual trial.

**Figure 3.**
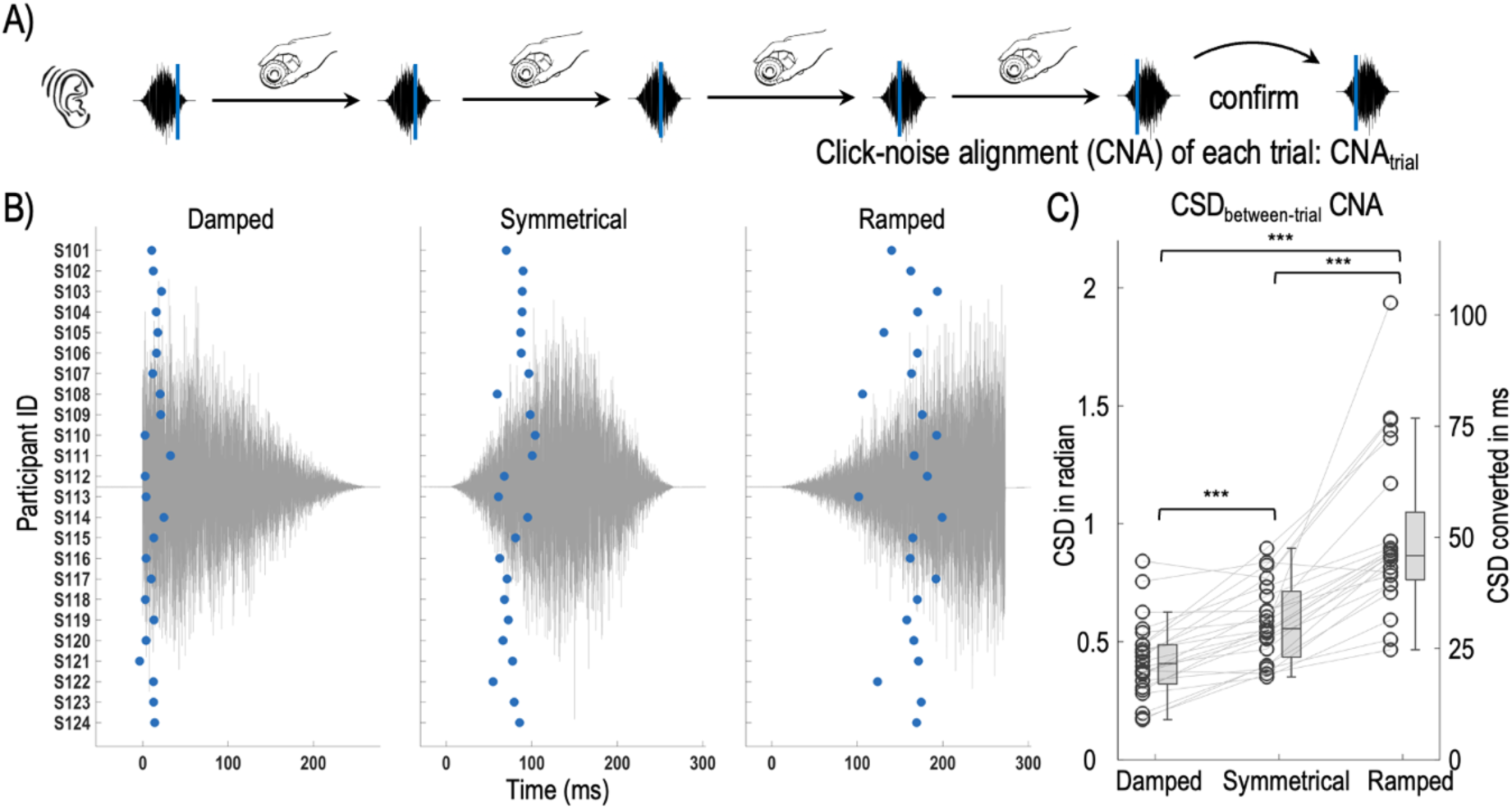
Participants’ between-trial performance in the sensory synchronization task. A) The click-noise alignment of each trial (CNA_trial_) was measured based on the final position of the tone click of trial that was confirmed by the participant when the tone click was perceived as in-sync with the noise stimulus. B) Average CNA of each stimulus envelope condition for individual participants (left: Damped stimuli; middle: Symmetrical stimuli; right: Ramped stimuli). In each graph, the grey curve indicates the wave form of an exemplar stimulus with the corresponding stimulus envelope. X-axis presents time, with zero indicating the onset of amplitude rise. The blue circles indicate the condition-average of CNA (CNA_condition_) of each of the 24 participants. Y-axis shows participant ID, which are common across the three graphs. C) Between-trial variation of CNA of the three stimulus envelope conditions (CSD_between-trial_ CNA) of the three stimulus envelope conditions. For each envelope condition, circles indicate the average CSD_between-trial_ CNA of individual participants. The levels of CSD are shown in both radians (y-axis on the left side) and ms (reversely converted from circular data) (y-axis on the right side). Asterisks indicate statistical significance of pair-wise comparisons among envelope conditions (***: *p* < .001).

**Figure 4.**
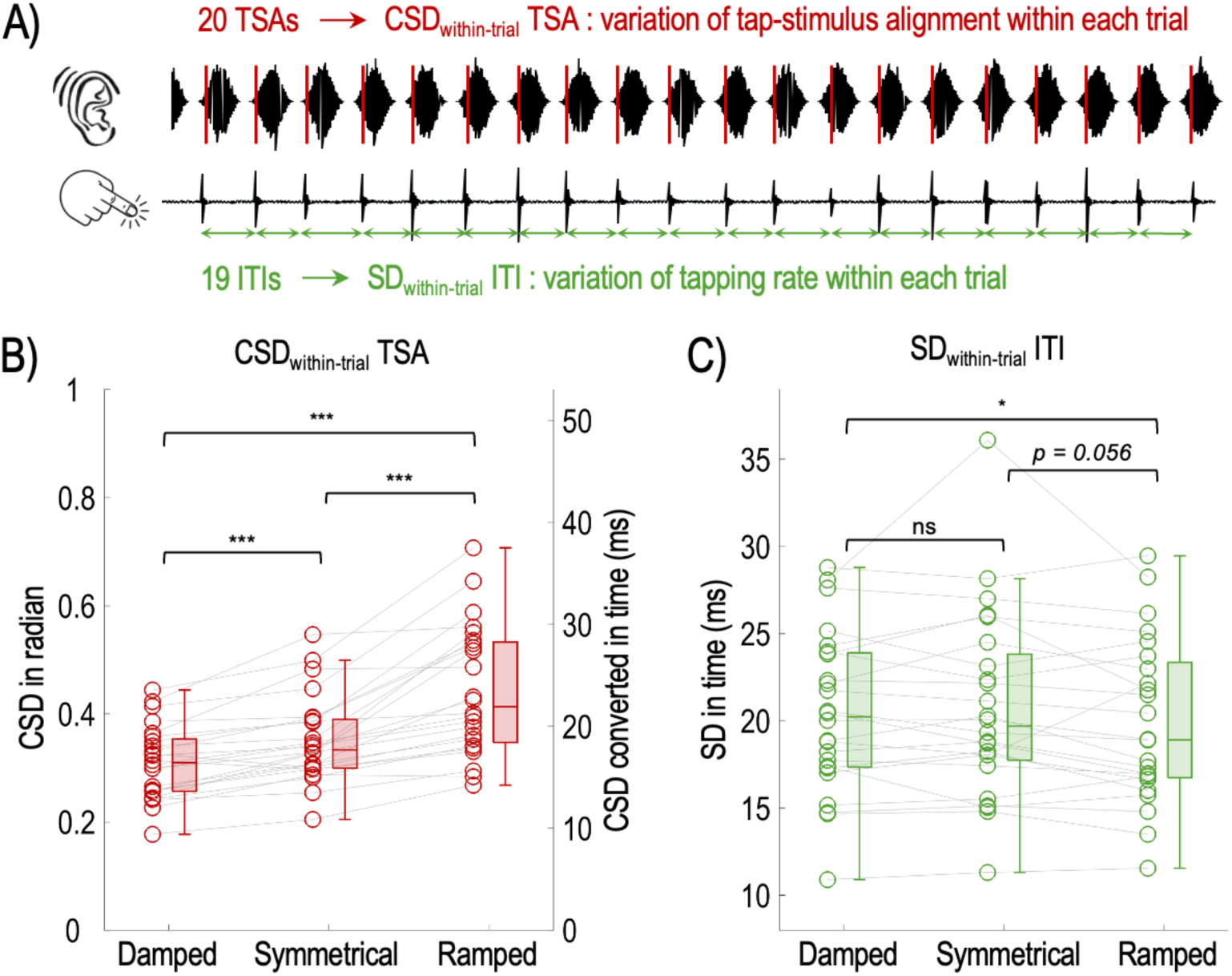
Participants’ within-trial tapping performances. A) Measurements of within-trial variation of tap-stimulus alignments (TSA) and inter-tap-interval (ITI). B) Within-trial variation of TSA of the three stimulus envelope conditions (CSD_within-trial_ TSA). For each envelope condition, circles indicate the average CSD_within-trial_ TSA of individual participants. The levels of CSD are shown in both radians (y-axis on the left side) and ms (reversely converted from circular data) (y-axis on the right side). Asterisks indicate statistical significance of pair-wise comparisons among envelope conditions (*: *p* <.05; ***: *p* < .001). C) Within-trial variation of ITI of the three stimulus envelope conditions (SD_within-trial_ ITI). For each envelope condition, circles indicate the average SD_within-trial_ ITI of individual participants.

##### Statistical analysis

To examined the effect of stimulus envelope shape on the between-trial variability of participants’ TSAs, we used a non-parametric Friedman test with participants’ CSD_between-trial T_SA as the dependent variable and stimulus envelope as the within-participant factor (damped *vs.* symmetrical *vs.* ramped). In case of significant effect of stimulus envelope, we ran post-hoc comparisons with Wilcoxon signed-rank tests. We conducted the same analyses on participants’ within-trial variability of TSAs (CSD_within-trial T_SA) and of ITIs (SD_within-trial I_TI). See results for detailed descriptions of these analyses.

##### Removal of trials with outlier synchronization performances

All participants self-reported having achieved synchronized tapping for most trials across the three envelope conditions. Nevertheless, we ran a procedure for each participant’s data to detect and remove outlier trials in which the participant gave relatively inferior synchronization performance.

For each participant and envelope condition, we removed trials which, across the final 20 taps, exhibited exceptionally irregular tapping rates (ITIs) and/or tap-stimulus alignments with respect to other trials from the same condition. We conducted outlier removals for each envelope condition in order to avoid unfairly removing more trials from conditions which presumably elicited less consistent synchronized tapping than other conditions. Our goal was to achieve a more uniform trial cohort that reflect participants’ synchronization performances in each envelope condition.

For each condition, trials whose mean tapping rate (reflected in ITI_trial)_ exceeded the 99.7% percentile (equivalent to 3 times of the standard deviation of both sides) of the condition distribution (estimated as a normal distribution) were considered as outliers and removed. Trials whose within-trial ITI variation (SD_within-trial I_TI) was higher than the 99.7% percentile (right side) of the condition distribution (estimated as a Chi2 distribution) were considered as outliers and removed. Trials whose within-trial TSA variation (CSD_within-trial T_SA) was higher than the 99.7% percentile (right side) of the condition distribution (estimated as a Chi2 distribution) were considered as outliers and removed.

In summary, our procedure removed trials with exceptionally irregular tapping rates and/or tap-stimulus alignments, while preserving the effect of each envelope condition on both measurements. Our procedure removed an average of 5.83% of trials per participant.

#### Sensory synchronization task

##### Determine the click-noise alignment for each trial

For this task, the experimental control program recorded the temporal position of the tone click within last modulation window of each trial. This final temporal position was selected and confirmed by the participant when the tone click was perceived as in-sync with the AM noise sequence (see Figure 3A). For each trial, we calculated the delay (in ms) between the onset of final tone click and the onset of the corresponding modulation window, which indicated the *click-noise alignment* of the trial (CNA_trial)_. Same as for the tap-stimulus alignment (TSA), we converted time delays (ms) into angles (radian) within a circular space that covered the modulation window in order to more accurately estimate the average click-noise alignment for each trial.

##### Between-trial synchronization performance

We then computed two measurements in order to assess participants’ between-trial synchronization performance in the sensory synchronization task. First, we computed the average CNA across all the trials of each envelope condition (CNA_condition)_, which reflected each participant’s overall CNA locations for each envelope shape. To examine the level of consistency with which participants aligned the tone click to the noise stimuli across different trials of each envelope condition, we computed the circular standard deviation (Mardia, 1975) across CNAs from different trials of each envelope condition (CSD_between-trial C_NA). Consistent click-noise alignments between trials should lead to low CSD_between-trial C_NA for individual participants.

##### Statistical analyses

To examined the effect of stimulus envelope shape on the between-trial variability of participants’ CNAs, we used a non-parametric Friedman test with participants’ CSD_between-trial C_NA as the dependent variable and stimulus envelope as the within-participant factor (damped *vs.* symmetrical *vs.* ramped). In case of significant effect of stimulus envelope, we ran post-hoc comparisons with Wilcoxon signed-rank tests.

## Results

### Effect of envelope shape on participants’ average tap-stimulus synchronization

#Figure 2B shows each participant’s average tap-stimulus alignment for each stimulus envelope (TSA_condition)_. Note that all TSA-related analyses were conducted using circular data (in radians) (see Methods). To better illustrate the TSA locations with respect to different types of envelopes, we displayed participants’ TSA_condition a_s time delays (in ms) to the AM onset of noise stimuli in Figure 2B. Also, in our results we report the average and variability of TSA in both radian and ms (converted from the circular data). For the damped stimuli, participants’ average taps are generally aligned to stimulus onset with a negative asynchrony (<u>converted time delay</u>: mean = -10.70 ms, SD = 11.60 ms; circular <u>TSA_condition_</u>: mean = -0.20 radian, CSD = 0.22 radian). For the other two stimulus envelopes with more gradual amplitude onsets, participants’ average taps were distributed after the onset of the amplitude rise, yielding an average delay of 44.87 ms (SD = 14.35 ms) across participants for the symmetrical envelope (circular TSA_condition:_ mean = 0.85 radian, CSD = 0.27 radian) and an average delay of 115.01 ms (SD = 40.68 ms) for the ramped envelope (circular TSA_condition:_ mean = 2.17 radian, CSD = 0.77 radian).

We then examined the between-trial variability of participants’ TSAs to different stimulus envelopes (Figure 2C). We used a non-parametric Friedman test with participants’ CSD_between-trial T_SA as the dependent variable and stimulus envelope as the within-participant factor. The test showed a significant effect of stimulus envelope (Chi-2(2) = 46.08, *p* < .001). Post-hoc comparisons with Wilcoxon signed-rank tests revealed higher CSD_between-trial T_SA for the ramped envelope than for the symmetrical (Ramped - Symmetrical = 0.45 radian (23.78 ms); *z* = 4.29; *p* < .001) and for the damped conditions (Ramped - Damped = 0.57 radian (30.03 ms); *z* = 4.29; *p* < .001), as well as higher CSD_between-trial T_SA for the symmetrical condition than for the damped condition (Symmetrical - Damped = 0.12 radian (6.24 ms); *z* = 4.09; *p* < .001).

In summary, participants exhibited larger between-trial variability in their tap-stimulus alignments when synchronized to stimuli with more gradual onsets than to those with sharper onsets. This result aligns with findings from a previous study that employed sensorimotor synchronization tasks to musical and quasi-musical stimuli with different onset dynamics (Danielsen et al., 2019).

#### Effect of envelope shape on participants’ click-noise synchronization

Next, the analysis of the sensory synchronization task revealed similar between-trial performance as the motor synchronization task. First, participants’ average click-noise alignment (CNA_condition)_ for different types of envelope shapes showed similar distributional properties as their average tap-stimulus alignment (TSA_condition)_ for the same envelope condition (Figure 3B). For the damped stimuli, participants’ final click locations were aligned to stimulus onset with a positive asynchrony, contrasting with the negative asynchrony of the sensorimotor tapping task (<u>time delay</u>: mean = 12.3 ms, SD = 8.18 ms; <u>circular CNA_condition_</u>: mean = 0.23 radian, CSD = 0.15 radian). For the symmetrical stimuli, participants’ final click locations showed an average delay of 79.86 ms (SD = 13.98 ms) with respect to the onset of amplitude modulation (circular CNA_condition:_ mean = 1.51 radian, CSD = 0.26 radian). For the ramped stimuli, participants’ final click locations showed an average delay of 163.93 ms (SD = 24.94 ms) with respect to the onset of amplitude modulation (circular CNA_condition:_ mean = 3.09 radian, CSD = 0.47 radian).

Second, we examined the between-trial variability of participants’ CNAs for different stimulus envelopes. A Friedman test with participants’ CSD_between-trial C_NA as the dependent variable and stimulus envelope as the within-participant factor revealed a significant effect of stimulus envelope (Chi-2(2) = 42.25, *p* < .001). Post-hoc comparisons with Wilcoxon signed-rank tests revealed higher CSD_between-trial C_NA for the ramped envelope than for the symmetrical (Ramped - Symmetrical = 0.37 radian (19.61 ms); *z* = 4.26; *p* < .001) and for the damped conditions (Ramped - Damped = 0.53 radian (27.94 ms); *z* = 4.29; *p* < .001), as well as higher CSD_between-trial T_SA for the symmetrical condition than for the damped condition (Symmetrical - Damped = 0.16 radian (8.33 ms); *z* = 4.06; *p* < .001).

Finally, we compared participants’ average synchronization locations in their motor task and sensory task. For each participant, we calculated the angular difference between the average tap-stimulus alignment (TSA_condition)_ and the average click-stimulus alignment (CNA_condition)_ for each stimulus envelope. We referred to this measurement as motor-sensory asynchrony (MSA). All the three envelope shapes yielded a negative MSA across participants: damped stimuli (angular difference: mean = -0.43 radian; CSD = 0.22; converted time difference: mean = -23.04 ms; SD = 11.61 ms); symmetrical stimuli (angular difference: mean = -0.66 radian; CSD = 0.35 radian; converted time difference: mean = -34.94 ms; SD = 18.50 ms); ramped stimuli (angular difference: mean = -0.87 radian; CSD = 0.86 radian; converted time difference: mean = -46.36 ms; SD = 45.41 ms). This finding supported the presence of negative mean asynchrony between participants’ tap locations and the presumable locations of the acoustic landmarks to which participants synchronize their taps.

In summary, the results from both motor and sensory synchronization tasks converge to show more variable synchronization locations with gradual-onset stimuli than within the envelope of sharp-onset stimuli. This variability effect was mainly observed across different trials within individual participants, although more variable alignments to gradual-onset stimuli were also observable across different participants (Figure 2B, 3B). In the following analyses, we focus on participants’ motor synchronization performances on individual trials. The objective of these next analyses is to assess the impact of stimulus envelope shape on participants’ ability to generate and sustain temporally regular motor output that is synchronized to the input acoustic sequences.

#### Effect of envelope shape on participants’ continuous synchronization to acoustic input

We examined two characteristics of participants’ continuous motor output across the different taps in individual trials (see Methods and Materials for details): (1) the maintenance of tap-stimulus alignments (TSA) across different taps which was reflected in the variation of TSAs across taps of individual trials (CSD_within-trial T_SA) (Figure 4A); (2) the maintenance of tapping rate which was reflected in the variation of ITIs across different taps of individual trials (SD_within-trial I_TI) (Figure 4A).

We first examined the impact of envelope shape on participants’ TSAs within trials. A Friedman test with participants’ average CSD_within-trial T_SA as the dependent variable and stimulus envelope as the within-participant factor revealed a main effect of stimulus envelope (Chi-2(2) = 42.25, *p* < .001). Post-hoc comparisons with Wilcoxon signed-rank tests showed higher CSD_within-trial T_SA for the ramped envelope than for the symmetrical (Ramped - Symmetrical = 0.09 radian (4.84 ms); *z* = 4.26; *p* < .001) and for the damped conditions (Ramped - Damped = 0.13 radian (7.03 ms); *z* = 4.29; *p* < .001), as well as higher CSD_within-trial T_SA for the symmetrical condition than for the damped condition (Symmetrical - Damped = 0.04 radian (2.19 ms); *z* = 3.97; *p* < .001) (Figure 4B).

Next, we examined the impact of envelope shape on participants’ ITIs within trials. Overall, all the three envelope shapes yielded an average ITI close to 333 ms (Damped: mean = 333.26 ms, SD = 0.48 ms; Symmetrical: mean = 333.56 ms, SD = 0.58 ms; Ramped: mean = 334.11 ms, SD = 1.28 ms), indicating a tapping rate close to 3 Hz. To examine the impact of envelope shape on participants’ within-trial ITI variations, we conducted a Friedman test with SD_within-trial I_TI as the dependent variable and envelope shape as the within-participant factor. This analysis showed a marginally significant main effect of envelope shape (Chi-2(2) = 5.25, *p* = .072) (Figure 4C). Despite the lack of a significant main effect, we conducted comparisons among conditions to inspect potential trends. These comparisons revealed a significant difference between ramped and damped envelopes, but with the ramped envelope exhibiting lower within-trial ITI variations than the damped envelope (Ramped - Damped = -0.49 ms, *z* = -2.14, *p* = 0.03) (Figure 4C). There was also a marginally significant difference between ramped and symmetrical envelopes, also with the ramped envelope exhibiting lower within-trial ITI variations (Ramped - Symmetrical = -0.80 ms, *z* = -1.91, *p* = 0.056) (Figure 4C).

Our analyses reveal complex modulations of participants’ within-trial synchronized tapping performances by the shape of stimulus envelope. On the one hand, we demonstrated an effect of stimulus envelope on participants’ ability to maintain their tap alignment to the same part of the stimulus envelope within individual trials, with gradual-onset stimuli yielding larger TSA variations than sharp-onset stimuli. On the other hand, our analysis on within-trial ITIs showed no significant impact of stimulus envelope shape on the variations of ITIs across different taps from the same trial. The differential influence of envelope shape on within-trial TSA and ITI variations is intriguing, as one might expect a rather straightforward correspondence between the two measurements: more variable tap-stimulus alignment between consecutive taps should lead to more variable intervals between consecutive taps. Therefore, it is puzzling how participants exhibit larger difficulty in maintaining consistent tap-stimulus alignment in trials with the gradual-onset stimuli than with the sharp-onset stimuli, while being able to maintaining their inter-tap-intervals similarly consistently for both envelope types. To address this unexpected pattern, we conducted a more detailed examination on the trajectories of participants’ tap locations within trials.

#### Analyses of tap trajectory reveal stronger presence of TSA drift in synchronized tapping to ramped stimuli

Visual inspection revealed potential links between participants’ within-trial tap trajectories and the levels of ITI and TSA variations. Figure 5B shows the TSA trajectories across the 20 taps from three selected trials of a single participant (Participant 117) (Figure 5A). Based on the previous findings, we are particularly interested in exploring characteristics of participants’ tap trajectories that could result in different levels of TSA variations - while exerting little impact on ITI variation. Accordingly, the three selected trials illustrated in Figure 5B exhibit different levels of within-trial TSA variations while showing nearly identical within-trial ITI variations (Figure 5A). Visual inspection of their tap trajectories reveals that the increase of TSA variations coincided with the amount of TSA drifts during the course of the trial (Figure 5B). Meanwhile, the degree of TSA drifts did not impact the ITI variations of the three trials.

**Figure 5.**
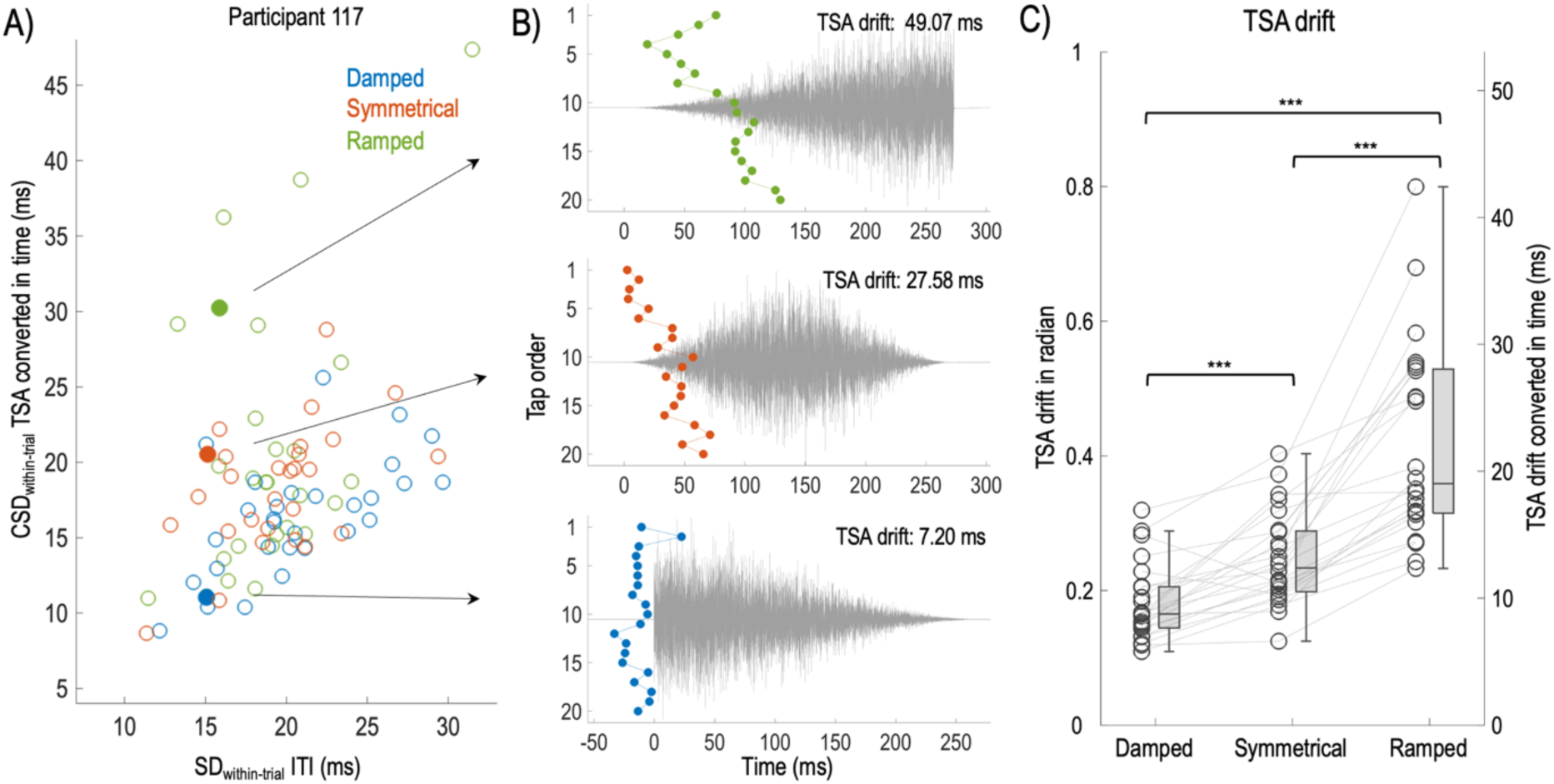
Analyses of tap trajectories within individual trials. A) Within-trial variations of tap-stimulus alignment (TSA) and inter-tap-interval (ITI) of trials from Participant 117. Filled circles indicate three trials that we selected to show their TSA trajectories across the 20 taps in Figure 5B (indicated by the arrows). These three trials exhibited different levels of within-trial TSA variations (CSD_within-trial_ TSA, y-axis) while showing similar within-trial ITI variations (SD_within-trial_ ITI, x-axis). B) Trajectories of tap-stimulus alignments across 20 taps of the three selected trials. In each graph, the x-axis presents time, with zero indicating the onset of the noise stimulus; y-axis present the order of the 20 taps from top to bottom; circles indicate the TSA location of each individual tap. C) Average TSA drift of the three stimulus conditions. For each envelope condition, circles indicate the average TSA drift of individual participants. The levels of TSA drift are shown in both radians (y-axis on the left side) and ms (reversely converted from circular data) (y-axis on the right side). Asterisks indicate statistical significance of pair-wise comparisons among envelope conditions (***: *p* < .001).

Based on these observations, we examined the impact of stimulus envelope on the degree of within-trial TSA drift. To quantify the degree of TSA drift in each trial, we computed the average TSA location across the first 10 taps and across the last 10 taps of the 20-tap analysis window for each trial. The absolute difference between the two average TSA locations thus, indicates the amount of TSA drift that takes place during the course of the trials. We then conducted a Friedman test on TSA drift with envelope shape as within-participant factor. The test revealed a main effect of envelope (Chi-2(2) = 44.33, *p* < .001). Post-hoc analyses revealed larger TSA drifts in participants’ taps to ramped stimuli than to the symmetrical (Ramped - Symmetrical = 0.17 radian (9.04 ms); *z* = 4.29; *p* < .001) and damped stimuli (Ramped - Damped = 0.24 radian (12.58 ms); *z* = 4.29; *p* < .001), as well as larger TSA drifts for the symmetrical stimuli than for damped stimuli (Symmetrical - Damped = 0.067 radian (3.54 ms); *z* = 3.77; *p* < .001) (Figure 5C).

In summary, our analysis of within-trial tap trajectories showed greater TSA drift when participants synchronously tap to stimuli with gradual-onset envelopes than to those with sharp-onset envelopes. While it is expected that larger TSA drifts leads to higher within-trial TSA variations, it did not seem to be impactful for the level of within-trial ITI variations.

#### A unified model to account for sensory and motor alignment with noise stimuli

Results from both motor and sensory synchronization tasks reveal that stimuli with gradual amplitude onsets lead to larger variability in participants’ alignment locations across different trials: tap-stimulus alignment (TSA) for the motor task and click-noise alignment (CNA) for the sensory task. This pattern indicates a relatively broad zone within the envelope of these stimuli that allows participants to align either their motor output or a sensory probe in order to achieve perceptual synchronization with the noise stimuli. In contrast, the small between-trial variability in TSA and CNA locations for sharp-onset stimuli indicates a narrow zone within these envelopes for achieving synchronization.

We outline a possible model which qualitatively accounts for this aspect of the alignment results from both tasks. The model characterizes a process which determines, for each individual trial, the location within the envelope of the noise stimuli which results in maximum synchrony between the noise stimuli and participants’ taps (in the motor task) or the tone click (in the sensory task). As a proof of concept, we use the synchronization process in the sensory task to illustrate the proposed mechanism, given that the computations of maximum synchrony between tone clicks and noise is restricted to the auditory domain and is more straightforward to implement.

The model is shown in Figure 6A. The core computational unit takes two inputs: the cochlear output of the modulated noise sequence (right section of diagram) and that of the tone-click sequence (left part of diagram). Since the tone clicks are at 1 kHz, we assume that the cochlear region at 1 kHz is relevant for the alignment task. The two cochlear envelopes at 1 kHz reflect instantaneous firing rate functions of the neuronal circuits that respond to the noise stimuli and tone clicks, respectively. In the computational realization of our model, maximum synchrony is determined by the sequential cascade of two neuronal circuits. First, the cochlear envelopes are processed by an *upward-level-crossing detector*, which produces a spike when the up-going envelope crosses a certain threshold. The timing of the spike depends on the level of the threshold. Second, a *coincidence detector* is modeled to be the Euclidean distance between the spike time from the noise envelope, which serves as the reference, and the spike time of the click envelope (which in the experiment can be continuously adjusted via a dial). Maximum synchrony is reached when the adjustment of click time leads to minimum distance with the noise time, which results in the optimal temporal alignment between the two auditory streams.

**Figure 6.**
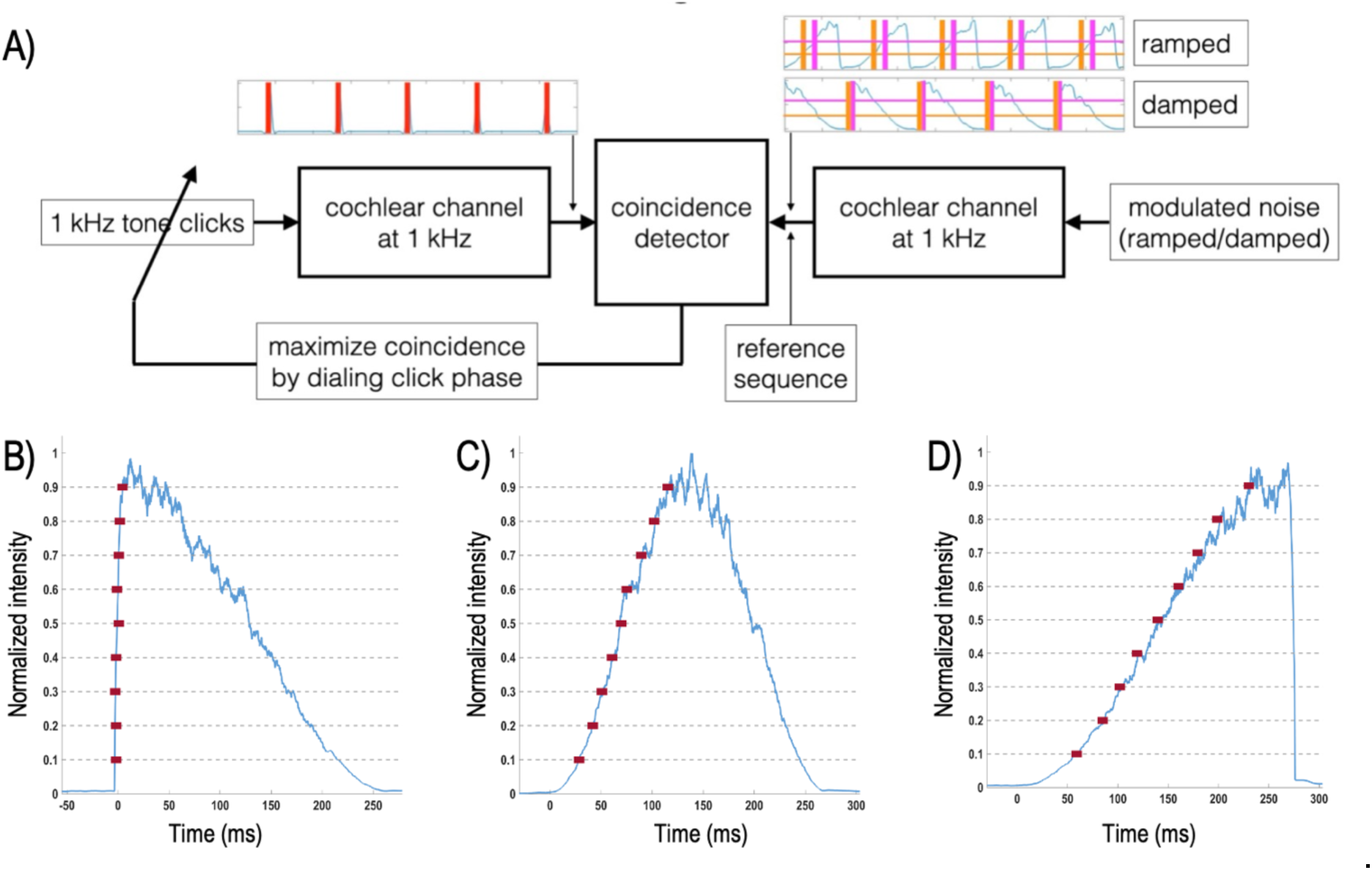
Model for the sensory synchronization task. A) Schematic presentation of the model. A coincidence detector provides the distance between reference spikes, driven by the modulated noise stimuli as well as the clicks. The search for best alignment is accomplished via dialing the click phase towards minimum distance. The neuronal circuit that generates spikes is modeled as an upward going level-crossing detector, and the parameter is the level-crossing threshold. The figure depicts spike sequences for two threshold values (in orange and in magenta), overlayed on top of the corresponding simulated cochlear envelopes, for ramped and damped stimuli. B-D) Locations of the best alignment as a function of threshold level of the upward-level-crossing-detector for damped stimuli (B), symmetrical stimuli (C) and ramped stimuli (D). The blue curves represent the amplitude envelope of the 1kHz cochlear channel of the modulated noise stimuli. Dotted lines indicate the level of threshold. Red squares indicate the locations of the optimal synchronization positions that correspond to the intersection between each threshold level and noise envelope.

Given the transient, sharp nature of the tone clicks, which should result in constant spike times with respect to their onset, the position of the optimal alignment ultimately depends on the spike time generated from the envelope of the modulated noise stimuli. Our model considers two sources of variability that affect the spike time of the noise stimuli. The first one is caused by biophysical uncertainties in the neuronal circuit of the upward-level-crossing detector, which changes the level of the threshold from one trial to another. The second one is caused by the random amplitude fluctuations in the envelope of the noise stimuli given the stochasticity of the broadband noise used in our study. Both sources of variability affect the location where the envelope of noise stimuli meets the threshold level, and hence, the resulting spike time.

Outcomes of the model show that the spike times of gradual-onset stimuli are sensitive to the level of threshold (Figure 6C, D), while those of sharp-onset stimuli are not (Figure 6B). As shown in the figures, fluctuations of the threshold level within a fixed range lead to wider distributions of the spike time from the envelopes of gradual-onset stimuli than from those of sharp-onset stimuli, which consequently results in wider distribution of optimal alignment locations for the former. This outcome is qualitatively in line with larger between-trial variability of click-noise alignments for gradual-onset stimuli than for sharp-onset stimuli observed in the sensory task (Figure 3), which suggests a reset of the biophysical threshold at the start of each trial.

The model also shows that the stochasticity of the broadband noise generates additional variation in spike time for each threshold level. This variation is also larger for gradual-onset stimuli than for sharp-onset stimuli. Under the assumption of a fixed threshold level for each trial, the additional variation in spike time should manifest across individual cycles of noise stimuli. Given that we only recorded the final alignment location of the tone click of each trial, such variation cannot be reflected in our data. Meanwhile, one could speculate that these additional variations may affect participants’ certainty level in selecting the best alignment location, with less certainty for gradual-onset stimuli than for sharp-onset stimuli.

Finally, although this model is based on sensory synchronization task, its architecture can also be generalized to qualitatively account for the finger tapping data. For that task, the act of tapping replaces the tone dialing: the subject aims at minimizing the distance between the spike time from the noise envelope and the perceived timing of their own taps (somatosensory input). The determination of the spike time of noise stimuli can be implemented in the same way as the sensory synchronization model, with the same two sources of variation affecting the spike time. Meanwhile, the determination of spike time for tapping would mainly involve sensory processing in the somatosensory modality. Furthermore, given the sequential nature of continuous tapping, participants should also be able to make use of their preceding tapping intervals, in addition to the spike time of the noise stimuli, to adjust the timing of their following taps (Jacoby & Repp, 2012).

## Discussion

We investigated how human listeners’ behavioral synchronization to periodic acoustic input is influenced by the shape of stimulus’ amplitude envelope. Our results revealed a complex effect. On one hand, we showed that the envelope shape strongly influenced the level of consistency with which align their behavioral output to the stimulus envelope. This effect was robustly observed in both the motor and the sensory synchronization paradigms. On the other hand, the envelope shape did not show significant impact on participants’ ability to maintain their tapping rate at the modulation rate of stimulus sequence. These findings suggest that the underlying processes for the maintenance of tap-stimulus alignment and tapping rate exhibit different levels of reliance on the saliency of acoustic landmarks in the acoustic signal.

### Synchronous tracking of acoustic events relies on the saliency of acoustic input

Our results align with previous findings showing different neural responses related to event- and rate-tracking during exposure to rhythmic acoustic input. For instance, studies using EEG or MEG recordings commonly report stronger auditory ERPs following stimuli with sharp onsets (Doelling et al., 2019; Irsik et al., 2021; Thomson et al., 2009). Intracranial recordings further reveal specific neural populations within the auditory cortex of human and non-human primates whose firing rate is specifically modulated by the sharpness of stimulus onset (Liu & Wang, 2022; Oganian & Chang, 2019).

Given that evoked responses are ‘temporally focal’ to their sensory triggers, strong responses following sharp acoustic onsets should efficiently convey the timing of these onsets and facilitate the estimation of the timing for the upcoming onsets. The precise timing of salient acoustic events in a periodic sequence would, therefore, be informative for the motor system to sustain consistent alignment of the motor output to individual events within the acoustic input.

In the absence of salient acoustic landmarks, stimuli with gradual amplitude onsets have been shown to elicit weak evoked responses from human auditory cortex (Doelling et al., 2019; Irsik et al., 2021; Thomson et al., 2009) and lower firing rates in onset-sensitive neurons from human and non-human primates (Liu & Wang, 2022; Oganian & Chang, 2019). For instance, Liu and Wang (2022) showed that the spiking of onset-sensitive neurons in marmoset auditory cortex is temporally more scattered when the animals were exposed to stimuli with ramped amplitude onsets. Diminished evoked responses indicate weaker timing estimations of potential landmarks that can be extracted from gradual-onset envelopes. If participants rely on these landmarks to guide the timing of their tapping, they are expected to show less consistent tapping alignments to stimulus envelope. Our finding is in line with this interpretation, and suggests a functional link between auditory evoked responses and temporal tracking of repeated acoustic landmarks in an external signal.

### Successful rate tracking without salient acoustic landmarks

When listeners can clearly identify an acoustic landmark that is repeated in the continuous stimulus envelope, the rate of amplitude modulation can be deduced by computing the intervals between successive landmarks. In other words, given that salient acoustic landmarks lead to consistent alignments between individual taps and the corresponding landmarks, these consistent alignments should straightforwardly result in consistent tapping rate. While our results with sharp-onset stimuli confirm this expectation, we also demonstrate that participants can equally efficiently track the repetition rate from sequences with gradual-onset stimuli and match the rate of their motor output to it. In particular, comparable rate tracking performance with gradual-onset stimuli and sharp-onset stimuli was accompanied by less consistent tap alignments to the envelopes of gradual-onset stimuli than to those of sharp-onset stimuli. These observations indicate that tracking of temporal periodicity does not require precise timing processing of local acoustic landmarks from the repeated stimulus envelope.

Indeed, it has been demonstrated that fluctuations of neural activity in human auditory cortex can synchronize (or be entrained) to temporal regularities in external acoustic stimuli, such that the entrained neural activity fluctuates at the same rate as the external stimulation (Lakatos et al., 2019). These cortical signals have been proposed to be underpinned by oscillatory fluctuations of local neuronal excitability that manifest intrinsically in the auditory cortex (Buzsáki & Draguhn, 2004; Lakatos et al., 2005). Synchronization of these types of neural activity to external stimulation thus involves adjusting the period of intrinsic neural oscillations to match the modulation rate of the acoustic input (Poeppel & Teng, 2020; Thut et al., 2011). While the underlying mechanism for these adjustments is debated (Doelling & Assaneo, 2021), it is less controversial that entrainment of slow cortical activities to the rate of external stimulation can be achieved using temporal sequences that contain salient acoustic events (Lakatos et al., 2013) as well as those with gradually changing amplitude (Henry et al., 2014; Irsik et al., 2021).

Integrating findings on neural entrainment with our results, one can speculate that, in the absence of salient acoustic landmarks, the rate of the periodic noise sequence is extracted via entrainment of slow oscillatory activity in auditory cortex and relayed to the motor system for generating a rhythmic motor sequence at the entrained rate. This processing scheme is supported by previous findings on rhythmic coupling between auditory and motor areas (Assaneo & Poeppel, 2018; Poeppel & Assaneo, 2020).

### Corrective processes during synchronous tapping

Our study provides insights into the higher-order components of sensorimotor synchronization. Previous work has established two corrective processes with which participants continuously regulate the timing of their taps: (i) *phase correction*, which aims at minimizing the temporal mis-alignment between taps and a sensory reference; and (ii) *period correction*, which aims at maintaining the intervals between successive taps as close to a target duration as possible (Jacoby & Repp, 2012; Repp, 2005). It has been shown that, while both processes manifest during synchronized tapping, phase correction is the more prominent process in determining the timing of taps when participants tap along periodic sequences of metronome clicks with salient, sharp onsets (Repp, 2005).

Using stimuli with gradual onsets, we introduced a way to manipulate the strength of phase correction. Specifically, we revealed that the less consistent tap-stimulus alignment is caused by larger degrees of slow alignment drifts, which have been previously observed when participants try to sustain periodic tapping without external stimuli (i.e., self-paced tapping). The degree of slow drifts have thus been proposed to reflect the strength of period correction in the regulation of tap timing (Madison, 2001). Compared to previous studies, which focused on cases with either strong reliance on phase correction (synchronized tapping to metronome clicks) or dominant reliance on period correction (self-paced tapping), our study offers a way to manipulate the relative strengths of the two processes.

### Focal synchronization spots within a broad zone of the stimulus envelope

Our findings also complement previous research on the perceptual center of acoustic stimuli. This research mainly focuses on the location of a specific zone within the stimulus envelope that allows listeners to align their synchronous output. We showed that the shape of stimulus envelope modulates not only the location of participants’ alignment zones but also the broadness of those alignment zones. Indeed, larger between-trial variabilities of TSA and CNA for gradual-onset stimuli suggest wider synchronization zones in the envelopes of these stimuli compared to those of sharp-onset stimuli. These results confirm previous findings that showed an increase of the size of synchronization zones for musical and quasi-musical stimuli with gradual amplitude rises (Danielsen et al., 2019).

Interestingly, while the envelope of gradual-onset stimuli provided a relatively large zone that is viable for participants to synchronize their taps to, listeners can align their taps tightly around a specific spot within that zone in each trial. This phenomenon is more pronounced in the sensory task, where on each trial participants could freely explore various locations to align the tone click and the noise stimulus before explicitly choosing one optimal location that gave the percept of maximum synchrony between the two stimuli. It remains unclear what mechanism accounts for the variation of the optimal synchronization spot across different trials. In our hypothesized auditory model, we propose that this variation primarily arises from biophysical fluctuations within the neuronal circuit that detects the amplitude onset of auditory input, which may also underlie the scattered spiking time of auditory neurons following ramped sensory input (Liu & Wang, 2022).

## Conclusion

Our behavioral assessment of synchronization to external auditory rhythmicity highlights distinct processes for tracking individual acoustic events and the rate of periodicity in continuous signals. The data provide a meaningful link between neurophysiological responses to envelope characteristics of auditory input and the cognitive aspects in humans’ behavioral synchronization to sensory rhythmicity.

## Acknowledgements

We would like to thank Julia Guldan, Freya Materne, and Claudia Lehr for their assistance with data collection as well as Cornelius Abel and Patrick Ulrich for technical support. We are also grateful to Xiangbin Teng for thoughtful comments on various aspects of the study. This work was supported by the Max Planck Society and the Ernst Struengmann Institute for Neuroscience.

